# SARS-CoV-2 mRNA Vaccine Development Enabled by Prototype Pathogen Preparedness

**DOI:** 10.1101/2020.06.11.145920

**Authors:** Kizzmekia S. Corbett, Darin Edwards, Sarah R. Leist, Olubukola M. Abiona, Seyhan Boyoglu-Barnum, Rebecca A. Gillespie, Sunny Himansu, Alexandra Schäfer, Cynthia T. Ziwawo, Anthony T. DiPiazza, Kenneth H. Dinnon, Sayda M. Elbashir, Christine A. Shaw, Angela Woods, Ethan J. Fritch, David R. Martinez, Kevin W. Bock, Mahnaz Minai, Bianca M. Nagata, Geoffrey B. Hutchinson, Kapil Bahl, Dario Garcia-Dominguez, LingZhi Ma, Isabella Renzi, Wing-Pui Kong, Stephen D. Schmidt, Lingshu Wang, Yi Zhang, Laura J. Stevens, Emily Phung, Lauren A. Chang, Rebecca J. Loomis, Nedim Emil Altaras, Elisabeth Narayanan, Mihir Metkar, Vlad Presnyak, Catherine Liu, Mark K. Louder, Wei Shi, Kwanyee Leung, Eun Sung Yang, Ande West, Kendra L. Gully, Nianshuang Wang, Daniel Wrapp, Nicole A. Doria-Rose, Guillaume Stewart-Jones, Hamilton Bennett, Martha C. Nason, Tracy J. Ruckwardt, Jason S. McLellan, Mark R. Denison, James D. Chappell, Ian N. Moore, Kaitlyn M. Morabito, John R. Mascola, Ralph S. Baric, Andrea Carfi, Barney S. Graham

**Author notes:** Authors have equal contribution to this study (Kizzmekia S. Corbett, Darin Edwards, Sarah R. Leist).

## Abstract

A SARS-CoV-2 vaccine is needed to control the global COVID-19 public health crisis. Atomic-level structures directed the application of prefusion-stabilizing mutations that improved expression and immunogenicity of betacoronavirus spike proteins. Using this established immunogen design, the release of SARS-CoV-2 sequences triggered immediate rapid manufacturing of an mRNA vaccine expressing the prefusion-stabilized SARS-CoV-2 spike trimer (mRNA-1273). Here, we show that mRNA-1273 induces both potent neutralizing antibody and CD8 T cell responses and protects against SARS-CoV-2 infection in lungs and noses of mice without evidence of immunopathology. mRNA-1273 is currently in a Phase 2 clinical trial with a trajectory towards Phase 3 efficacy evaluation.

---

Since its emergence in December 2019, severe acute respiratory syndrome coronavirus 2 (SARS-CoV-2) has accounted for over 7 million cases of Coronavirus Disease 2019 (COVID-19) worldwide in less than 7 months^1^. SARS-CoV-2 is the third novel betacoronavirus in the last 20 years to cause substantial human disease; however, unlike its predecessors SARS-CoV and MERS-CoV, SARS-CoV-2 transmits efficiently from person-to-person. In absence of a vaccine, public health measures such as quarantining newly diagnosed cases, contact tracing, and mandating face masks and physical distancing have been instated to reduce transmission^2^. It is estimated that until 60-70% population immunity is established, it is unlikely for COVID-19 to be controlled well enough to resume normal activities. If immunity remains solely dependent on infection, even at a 1% mortality rate, >40 million people could succumb to COVID-19 globally^3^. Therefore, rapid development of vaccines against SARS-CoV-2 is critical for changing the global dynamic of this virus.

The spike (S) protein, a class I fusion glycoprotein analogous to influenza hemagglutinin (HA), respiratory syncytial virus (RSV) fusion glycoprotein (F), and human immunodeficiency virus (HIV) gp160 (Env), is the major surface protein on the CoV virion and the primary target for neutralizing antibodies. S proteins undergo dramatic structural rearrangement to fuse virus and host cell membranes, allowing delivery of the viral genome into target cells. We previously showed that prefusion-stabilized protein immunogens that preserve neutralization-sensitive epitopes are an effective vaccine strategy for enveloped viruses, such as RSV^4–8^. Subsequently, we identified 2 proline substitutions (2P) at the apex of the central helix and heptad repeat 1 that effectively stabilized MERS-CoV, SARS-CoV and HCoV-HKU1 S proteins in the prefusion conformation^9–11^. Similar to other prefusion-stabilized fusion proteins, MERS S-2P protein is more immunogenic at lower doses than wild-type S protein^11^. The 2P has been widely transferrable to other beta-CoV spike proteins, suggesting a generalizable approach for designing stabilized prefusion beta-CoV S vaccine antigens. This is fundamental to the prototype pathogen approach for pandemic preparedness^12,13^.

Coronaviruses have long been predicted to have a high likelihood of spill over into humans and cause future pandemics^14,15^. As part of our pandemic preparedness efforts, we have studied MERS-CoV as prototype pathogen for betacoronaviruses to optimize vaccine design, to dissect the humoral immune response to vaccination, and identify mechanisms and correlates of protection. Achieving an effective and rapid vaccine response to a newly emerging virus requires the precision afforded by structure-based antigen design but also a manufacturing platform to shorten time to product availability. Producing cell lines and clinical grade subunit protein typically takes more than 1 year, while manufacturing nucleic acid vaccines can be done in a matter of weeks^16,17^. In addition to advantages in manufacturing speed, mRNA vaccines are potently immunogenic and elicit both humoral and cellular immunity^18–20^. Therefore, we evaluated mRNA formulated in lipid nanoparticles (mRNA/LNP) as a delivery vehicle for the MERS S-2P and found that transmembrane-anchored MERS S-2P mRNA elicited better neutralizing antibody responses than secreted MERS S-2P **(Extended Data Fig. 1a)**. Additionally, consistent with protein immunogens, MERS S-2P mRNA was more immunogenic than MERS wild-type S mRNA **(Extended Data Fig. 1b)**. Immunization with MERS S-2P mRNA/LNP elicited potent neutralizing activity down to a 0.1 μg dose and protected hDPP4 transgenic (288/330^+/+21^) mice against lethal MERS-CoV challenge in a dose-dependent manner, establishing proof-of-concept that mRNA expressing the stabilized S-2P protein is protective. Notably, the sub-protective 0.01 μg dose of MERS S-2P mRNA did not cause exaggerated disease following MERS-CoV infection, but instead resulted in partial protection against weight loss followed by full recovery without evidence of enhanced illness **(Fig. 1)**.

**Figure 1.**
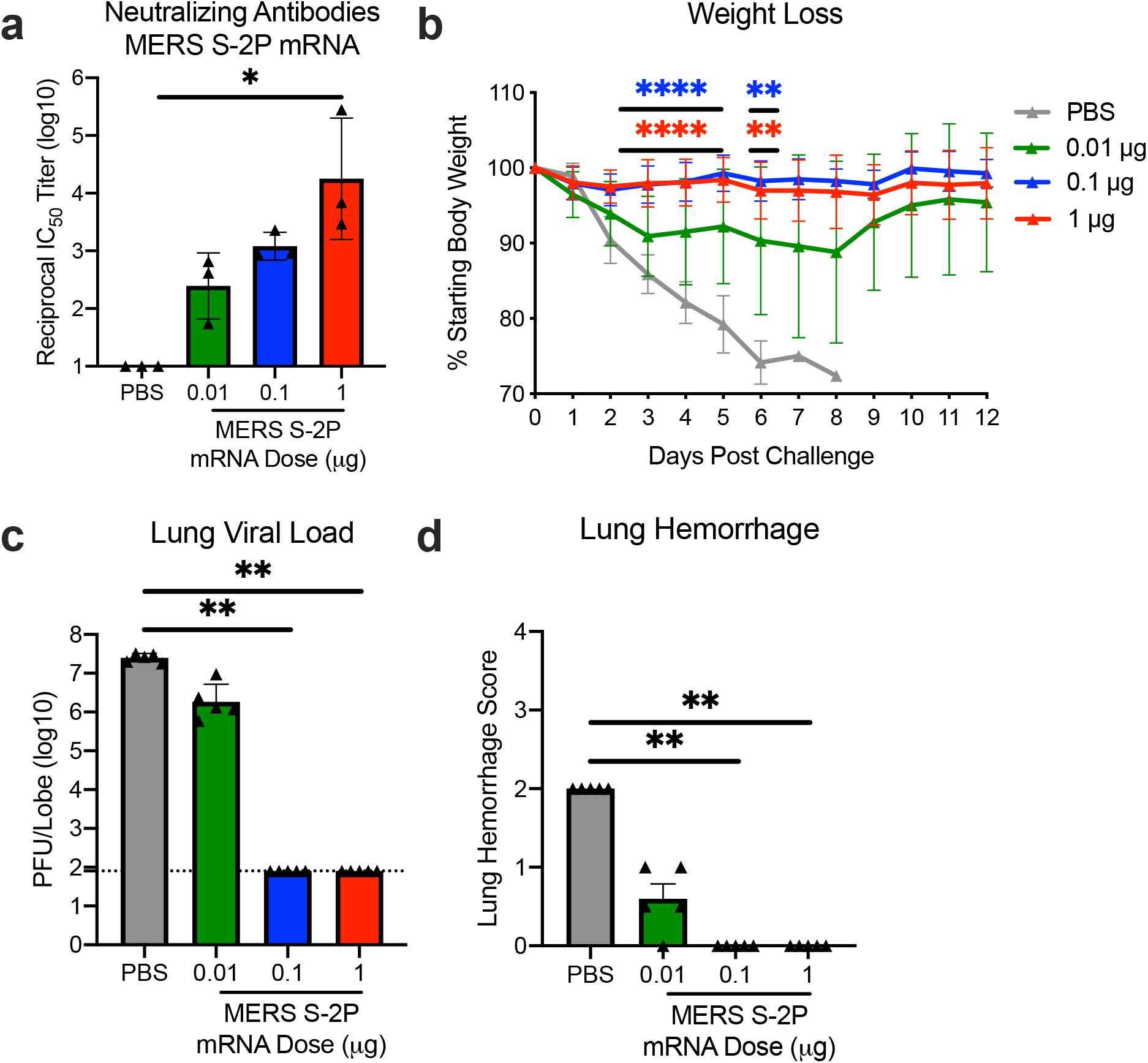
MERS-CoV S-2P mRNA protects mice from lethal challenge. 288/330^+/+^ mice were immunized at weeks 0 and 3 with 0.01 (green), 0.1 (blue), or 1 μg (red) of MERS-CoV S-2P mRNA. Mock-immunized mice were immunized with PBS (gray). Two weeks post-boost, sera were collected from 3 mice per group and assessed for neutralizing antibodies against MERS m35c4 pseudovirus (a). Four weeks post-boost, 12 mice per group were challenged with a lethal dose of mouse-adapted MERS-CoV (m35c4). Following challenge, mice were monitored for weight loss (b). Two days post-challenge, at peak viral load, lung viral titers (c) and hemorrhage (0 = no hemorrhage, 4 = severe hemorrhage in all lobes) (d) were assessed from 5 animals per group. Dotted line = assay limit of detection. (a, c-d) All dose levels were compared. (b) For weight loss, all comparisons are against PBS-immunized mice.

In early January 2020, a novel CoV (nCoV) was identified as the cause of a respiratory virus outbreak occurring in Wuhan, China. Within 24 hours of the release of the SARS-CoV-2 isolate sequences (then known as “2019-nCoV”) on January 10^th^, the 2P mutations were substituted into S positions aa986 and 987 to produce prefusion-stabilized SARS-CoV-2 S (S-2P) protein for structural analysis^22^ and serological assay development^23,24^ *in silico* without additional experimental validation. Within 5 days of sequence release, current Good Manufacturing Practice (cGMP) production of mRNA/LNP expressing the SARS-CoV-2 S-2P as a transmembrane-anchored protein with the native furin cleavage site (mRNA-1273) was initiated in parallel with preclinical evaluation. Remarkably, this led to the start of a first in human Phase 1 clinical trial on March 16, 2020, 66 days after the viral sequence was released, and a Phase 2 began 74 days later on May 29, 2020 **(Extended Data Fig. 2)**. Prior to vaccination of the first human subject, expression and antigenicity of the S-2P antigen delivered by mRNA was confirmed *in vitro* **(Extended Data Fig. 3),** and immunogenicity of mRNA-1273 was documented in several mouse strains. The results of those studies are detailed hereafter.

Immunogenicity was assessed in six-week old female BALB/cJ, C57BL/6J, and B6C3F1/J mice by immunizing intramuscularly (IM) twice with 0.01, 0.1, or 1 μg of mRNA-1273 at a 3-week interval. mRNA-1273 induced dose-dependent S-specific binding antibodies after prime and boost in all mouse strains **(Fig. 2a-c)**. Potent neutralizing activity was elicited by 1 μg of mRNA-1273, reaching 819, 89, and 1115 reciprocal IC_50_ geometric mean titer (GMT) for BALB/cJ, C57BL/6J, and B6C3F1/J mice, respectively **(Fig. 2d-f)**. These levels are similar to the neutralization activity achieved by immunizing with 1 μg of SAS-adjuvanted S-2P protein **(Extended Data Table 1)**. To further gauge immunogenicity across a wide dose range, BALB/cJ mice were immunized with 0.0025 – 20 μg of mRNA-1273 revealing a strong positive correlation between dose-dependent mRNA-1273-elicited binding and neutralizing antibody responses **(Extended Data Fig. 4)**. BALB/cJ mice that received a single dose of mRNA-1273 were evaluated in order to ascertain the utility for a one-dose vaccine regimen. S-binding antibodies were induced in mice immunized with one dose of 1 or 10 μg of mRNA-1273, and the 10 μg dose elicited neutralizing antibody activity that increased between week 2 and week 4, reaching 315 reciprocal IC_50_ GMT **(Extended Data Fig. 5a-b)**. These data demonstrate that mRNA expressing SARS-CoV-2 S-2P is a potent immunogen and neutralizing activity can be elicited with a single dose.

**Figure 2.**
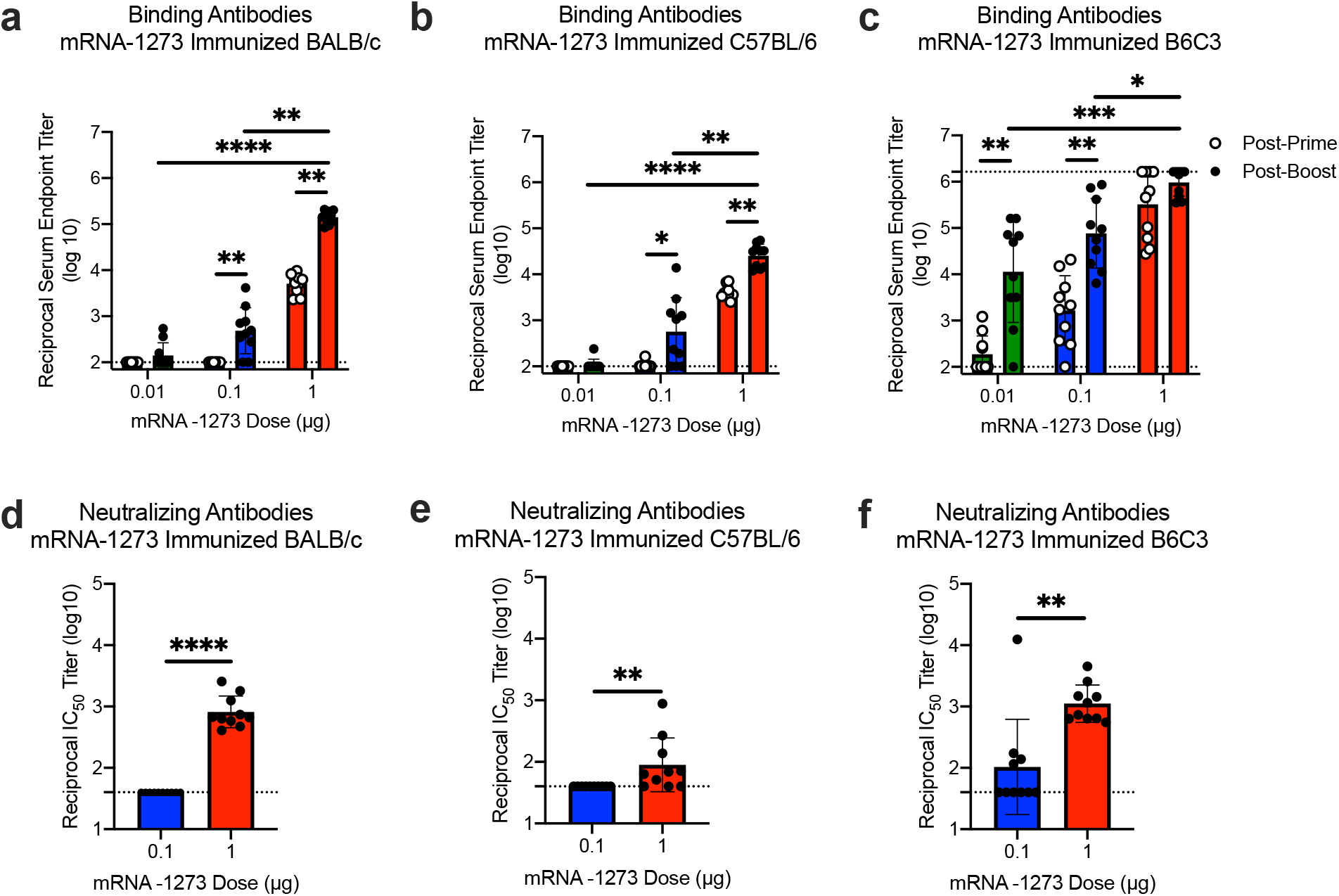
mRNA-1273 elicits robust binding and neutralizing antibody responses in multiple mouse strains. BALB/cJ (a, d), C57BL/6J (b, e), or B6C3F1/J (c, f) mice were immunized at weeks 0 and 3 weeks with 0.01 (green), 0.1 (blue), or 1 μg (red) of mRNA-1273. Sera were collected 2 weeks post-prime (open circles) and 2 weeks post-boost (closed circles) and assessed for SARS-CoV-2 S-specific IgG by ELISA (a-c), and, for post-boost sera, neutralizing antibodies against homotypic SARS-CoV-2 pseudovirus (d-f). Dotted line = assay limit of detection. (a-c) Timepoints were compared within each dose level, and doses were compared post-boost.

Next, we evaluated the balance of Th1 and Th2, because vaccine-associated enhanced respiratory disease (VAERD) has been associated with Th2-biased immune responses in children immunized with whole-inactivated virus vaccines against RSV and measles virus^25,26^. A similar phenomenon has also been reported in some animal models with whole-inactivated SARS-CoV vaccines^27^. Thus, we first compared levels of S-specific IgG2a/c and IgG1, which are surrogates of Th1 and Th2 responses respectively, elicited by mRNA-1273 to those elicited by SARS-CoV-2 S-2P protein adjuvanted with the TLR4-agonist Sigma Adjuvant System (SAS). Both immunogens elicited IgG2a and IgG1 subclass S-binding antibodies, indicating a balanced Th1/Th2 response **(Fig. 3a-c; Extended Data Fig. 6)**. The S-specific IgG subclass profile following a single dose of mRNA-1273 (**Extended Data Fig. 5c)** was similar to that observed following two doses. In contrast, Th2-biased antibodies with lower IgG2a/IgG1 subclass response ratios were observed in mice immunized with SARS-CoV-2 S protein formulated in alum **(Extended Data Fig. 7a-b)**. Following re-stimulation with peptide pools (S1 and S2) corresponding to the S protein, splenocytes from mRNA-1273-immunized mice secreted more IFN-γ than IL-4, IL-5, or IL-13 whereas SARS-CoV-2 S protein with alum induced Th2-skewed cytokine secretion **(Extended Data Fig. 7c-d)**. 7 weeks post-boost, we also directly measured cytokine patterns in vaccine-induced memory T cells by intracellular cytokine staining (ICS); mRNA-1273-elicited CD4+ T cells re-stimulated with S1 or S2 peptide pools exhibited a Th1-dominant response, particularly at higher immunogen doses **(Fig. 3d-e)**. Furthermore, 1 μg of mRNA-1273 induced a robust CD8+ T cell response to the S1 peptide pool **(Fig. 3f-g)**. The Ig subclass and T cell cytokine data together demonstrate that immunization with mRNA-1273 elicits a balanced Th1/Th2 response in contrast to the Th2-biased response seen with S protein adjuvanted with alum, suggesting that mRNA vaccination avoids Th2-biased immune responses that have been linked to VAERD.

**Figure 3.**
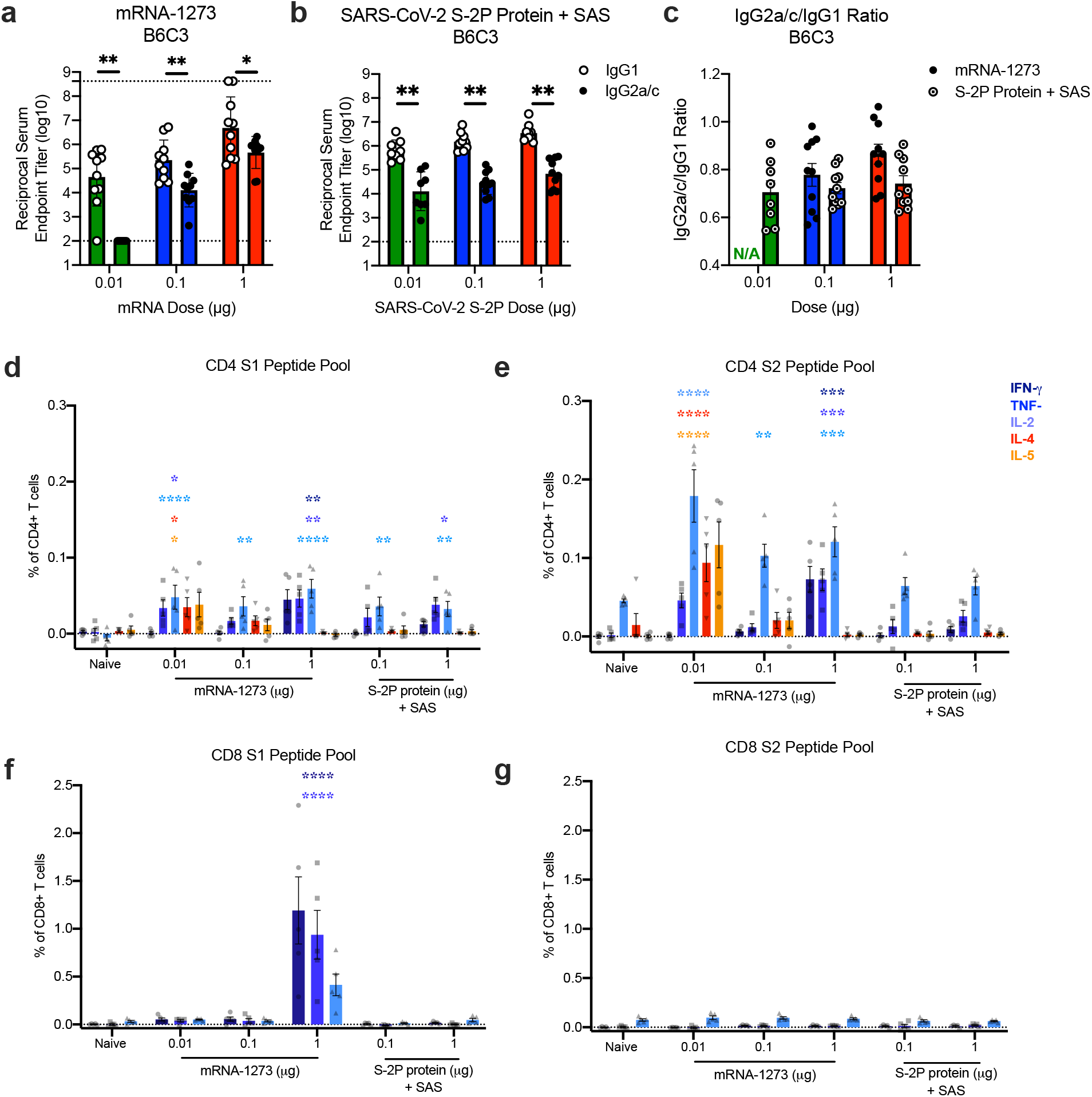
Immunizations with mRNA-1273 and S-2P protein, delivered with TLR4 agonist, elicit S-specific Th1-biased T cell responses. B6C3F1/J mice were immunized at weeks 0 and 3 with 0.01, 0.1, or 1 μg of mRNA-1273 or SAS-adjuvanted SARS-CoV-2 S-2P protein. Sera were collected 2 weeks post-boost and assessed by ELISA for SARS-CoV-2 S-specific IgG1 and IgG2a/c. Endpoint titers (a-b) and endpoint titer ratios of IgG2a/c to IgG1 (c) were calculated. For mice for which endpoint titers did not reach the lower limit of detection (dotted line), ratios were not calculated (N/A). (d-g) Seven weeks post-boost, splenocytes were isolated from 5 mice per group and re-stimulated with no peptides or pools of overlapping peptides from SARS-CoV-2 S protein in the presence of a protein transport inhibitor cocktail. After 6 hours, intracellular cytokine staining (ICS) was performed to quantify CD4+ and CD8+ T cell responses. Cytokine expression in the presence of no peptides was considered background and subtracted from the responses measured from the S1 and S2 peptide pools for each individual mouse. (d-e) CD4+ T cells expressing IFN-γ, TNFα, IL-2, IL-4 and IL-5 in response to the S1 (d) and S2 (e) peptide pools. (f-g) CD8+ T cells expressing IFN-γ, TNF-α, and IL-2 in response to the S1 (f) and S2 (g) peptide pools. IgG1 and IgG2a/c (a-b) and immunogens (c) were compared at each dose level. (d-g) For each cytokine, all comparisons were compared to naïve mice.

Protective immunity was assessed in young adult BALB/cJ mice challenged with mouse-adapted (MA) SARS-CoV-2 that exhibits viral replication localized to lungs and nasal turbinates^28^. BALB/cJ mice that received two 1 μg doses of mRNA-1273 were completely protected from viral replication in lungs after challenge at a 5- **(Fig. 4a)** or 13-week intervals following boost **(Extended Data Fig. 8a)**. mRNA-1273-induced immunity also rendered viral replication in nasal turbinates undetectable in 6 out of 7 mice **(Fig. 4b, Extended Data Fig. 8b)**. Efficacy of mRNA-1273 was dose-dependent, with two 0.1 μg mRNA-1273 doses reducing lung viral load by ~100-fold and two 0.01 μg mRNA-1273 doses reducing lung viral load by ~3-fold **(Fig. 4a)**. Of note, mice challenged 7 weeks after a single dose of 1 μg or 10 μg of mRNA-1273 were also completely protected against lung viral replication **(Fig. 4c)**. Challenging animals immunized with sub-protective doses provides an orthogonal assessment of safety signals, such as increased clinical illness or pathology. Similar to what was observed with MERS-CoV S-2P mRNA, mice immunized with sub-protective 0.1 and 0.01 μg mRNA-1273 doses showed no evidence of enhanced lung pathology or excessive mucus production **(Fig. 4d)**. In summary, mRNA-1273 is immunogenic, efficacious, and does not show evidence of promoting VAERD when given at sub-protective doses in mice.

**Figure 4.**
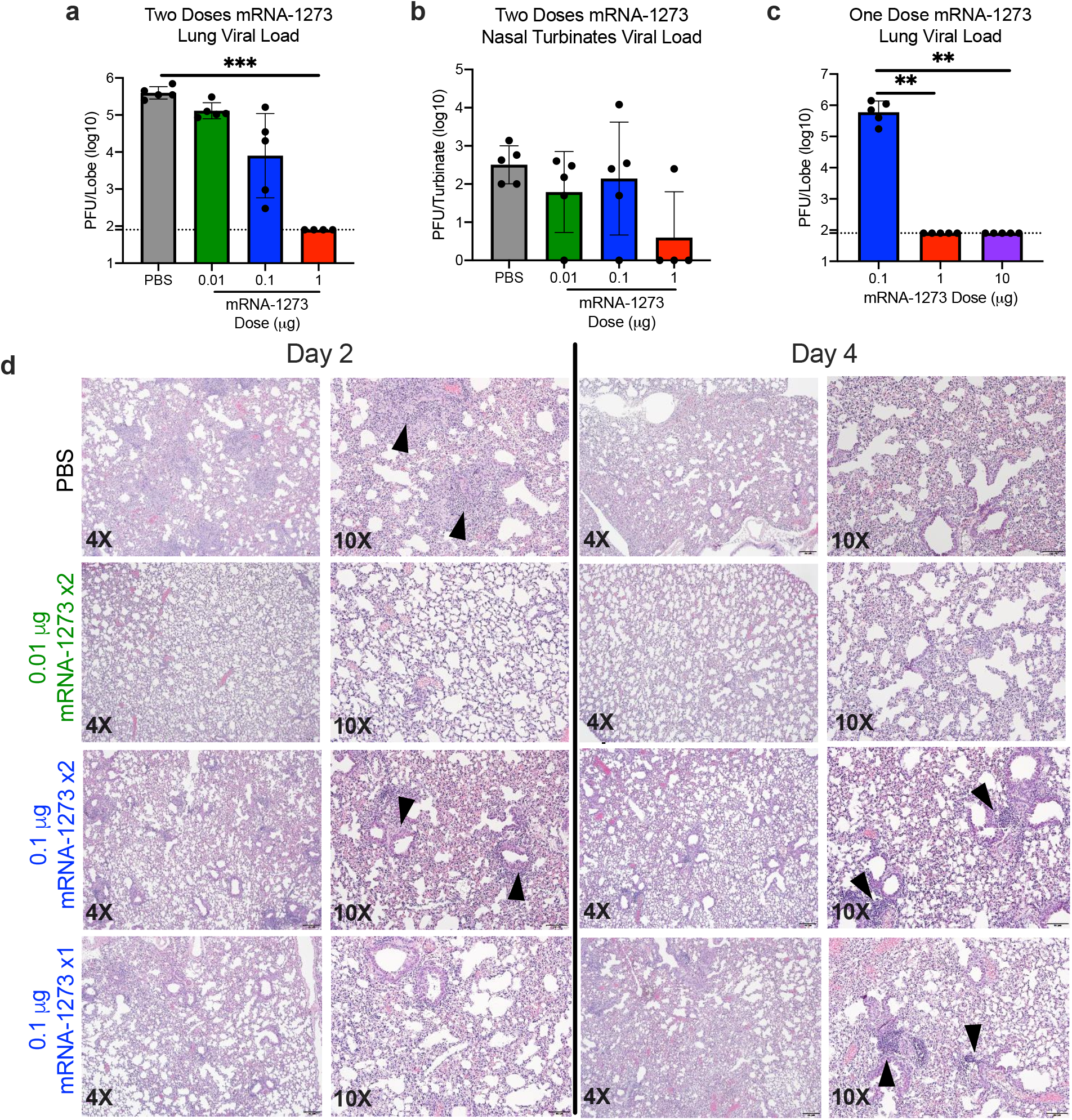
mRNA-1273 protects mice from upper and lower airway SARS-CoV-2 infection. (a-b) BALB/cJ mice were immunized at weeks 0 and 3 with 0.01 (green), 0.1 (blue), or 1 μg (red) of mRNA-1273. Mock-immunized mice were immunized with PBS x2. Five weeks post-boost, mice were challenged with mouse-adapted SARS-CoV-2. (c) BALB/cJ mice were also immunized with a single dose of 0.1 (blue),1 (red), or 10 (purple) μg of mRNA-1273 and challenged 7 weeks post-immunization. Two days post-challenge, at peak viral load, mouse lungs (a,c) and nasal turbinates (b) were harvested from 5 mice group for analysis of viral titers. Dotted line = assay limit of detection. (d) At day 2 and 4 post-challenge, lungs from 5 mice per group were fixed in 10% formalin, paraffin-embedded, cut in 5 μm sections, and stained with hematoxylin and eosin. Photomicrographs (4X and 10X) are representative of lung sections from groups of mice in which virus infection was detected. At day 2, lungs from mock-immunized mice demonstrated moderate to severe, predominantly neutrophilic, inflammation that was present within, and surrounding, small bronchioles (arrowheads); the surrounding alveolar capillaries were markedly expanded by infiltrating inflammatory cells. In the 0.01 μg two-dose group, inflammation was minimal to absent. In the 0.1 μg two-dose group, occasional areas of inflammation intimately associated with small airways (bronchioles) and their adjacent vasculature (arrowheads) were seen, primarily composed of neutrophils. In the single-dose 0.1 μg group, there were mild patchy expansion of the alveolar septae by mononuclear and polymorphonuclear cells. At day 4, lungs from mock-immunized mice exhibited moderate to marked expansion of the alveolar septae (interstitial pattern) with decreased prominence of the adjacent alveolar spaces. In the 0.01 μg two-dose group, inflammation was minimal to absent. Lungs in the 0.1 μg two-dose group showed mild, predominantly lymphocytic inflammation, intimately associated with bronchioles and adjacent vasculature (arrowheads). In the single-dose 0.1 μg group there was mild, predominantly lymphocytic, inflammation around bronchovascular bundles (arrowheads).

Here, we showed that 1 μg of mRNA-1273 was sufficient to induce robust neutralizing activity and CD8 T cell responses, balanced Th1/Th2 antibody isotype responses, and protection from viral replication for more than 3 months following a prime/boost regimen similar to that being tested in humans. Inclusion of lower sub-protective doses demonstrated the dose-dependence of antibody, Th1 CD4 T cell responses, and protection, suggesting immune correlates of protection can be further elucidated. A major goal of animal studies to support SARS-CoV-2 vaccine candidates through clinical trials is to not only prove elicitation of potent protective immune responses, but to show that sub-protective responses do not cause VAERD^3^. Sub-protective doses did not prime mice for enhanced immunopathology following challenge. Moreover, the induction of protective immunity following a single dose suggests that consideration could be given to administering one dose of this vaccine in the outbreak setting. These data, combined with immunogenicity data from nonhuman primates and subjects in early Phase 1 clinical trials, will be used to inform the dose and regimen of mRNA-1273 in advanced clinical efficacy trials.

The COVID-19 pandemic of 2020 is the Pathogen X event that has long been predicted^12,13^. Here, we provide a paradigm for rapid vaccine development. Structure-guided stabilization of the MERS-CoV S protein combined with a fast, scalable, and safe mRNA/LNP vaccine platform led to a generalizable beta-CoV vaccine solution that translated into a commercial mRNA vaccine delivery platform, paving the way for the rapid response to the COVID-19 outbreak. This is a demonstration of how the power of new technology-driven concepts like synthetic vaccinology facilitate a vaccine development program that can be initiated with pathogen sequences alone^11^. It is also a proof-of-concept for the prototype pathogen approach for pandemic preparedness and response that is predicated on identifying generalizable solutions for medical countermeasures within virus families or genera^12^. Even though the response to the COVID-19 pandemic is unprecedented in its speed and breadth, we envision a response that could be quicker. There are 24 other virus families known to infect humans, and with sustained investigation of those potential threats, we could be better prepared for future looming pandemics^13^.

## Acknowledgements

We thank Gabriela Alvarado, Karin Bok, Kevin Carlton, Masaru Kanekiyo, Robert Seder, and additional members of all included laboratories for critical discussions, advice, and review of the manuscript. We thank Judy Stein and Monique Young for technology transfer and administrative support, respectively. We thank members of the NIH NIAID VRC Translational Research Program for technical assistance with mouse experiments. This work was supported by the Intramural Research Program of the VRC and the Division of Intramural Research, NIAID, NIH (B.S.G) and NIH NIAID grant R01-AI127521 (J.S.M.). mRNA-1273 has been funded in part with Federal funds from the Department of Health and Human Services, Office of the Assistant Secretary for Preparedness and Response, Biomedical Advanced Research and Development Authority, under Contract 75A50120C00034. PRNT assays were funded under NIH Contract HHSN261200800001E Agreement 17×198 (to J.D.C.), furnished through Leidos Biomedical Research, Inc. MERS mRNA mouse challenge studies were funded under NIH Contract HHSN272201700036I Task Rrder No. 75N93019F00132 Requisition No. 5494549 (to R.B.). K.S.C.’s research fellowship was partially funded by the Undergraduate Scholarship Program, Office of Intramural Training and Education, Office of the Director, NIH. D.R.M. was funded by NIH NIAID grant T32-AI007151 and a Burroughs Wellcome Fund Postdoctoral Enrichment Program Award.

## Author Contributions

K.S.C., D.K.E., S.R.L., O.M.A, S.B.B., R.A.G., S.H., A.S., C.Z., A.T.D., K.H.D., S.E., C.A.S., A.W., E.J.F., D.R.M, K.W.B., M.M., B.M.N., G.B.H., K.B., D.G.D., L.M., I.R., W.P.K, S.S., L.W., Y.Z., J.C., L.S., L.A.C., E.P., R.J.L., N.E.A., E.N., M.M., V.P., C.L., M.K.L., W.S., K.G., K.L., E.S.Y., A.W., G.A., N.A.D.R., G.S.J., H.B., M.N., T.J.R., M.R.D., I.N.M., K.M.M., J.R.M., R.S.B., A.C., and B.S.G. designed, completed, and/or analyzed experiments. N.W., D.W., and J.S.M. contributed new reagents/analytic tools. K.S.C., K.M.M, and B.S.G. wrote the manuscript. All authors contributed to discussions in regard to and editing of the manuscript.

## Additional Information

Correspondence and requests for materials should be addressed to Barney S. Graham, and Andrea Carfi.

## Methods

### MERS-CoV S-2P and SARS-CoV-2 S-2P mRNA synthesis and lipid nanoparticle formulation

For each vaccine, T7 RNA polymerase-mediated transcription was used *in vitro* to synthesize the mRNA from a linearized DNA template, which flanked the immunogen open-reading frames with the 5′ and 3′ untranslated regions and a poly-A tail as described previously ^29^. mRNA was then purified, diluted in citrate buffer to the desired concentration and encapsulated into lipid nanoparticles (LNP) by ethanol drop nanoprecipitation. At molar ratio of 50:10:38.5:1.5 (ionizable lipid:DSPC:cholesterol:PEG-lipid), lipids were dissolved in ethanol and combined with a 6.25-mM sodium acetate buffer (pH 5) containing mRNA at a ratio of 3:1 (aqueous:ethanol). Formulations were dialyzed against phosphate-buffered saline (pH 7.4) for at least 18 hr, concentrated using Amicon ultracentrifugal filters (EMD Millipore), passed through a 0.22-μm filter and stored at −20°C until use. All formulations underwent quality control for particle size, RNA encapsulation, and endotoxin. LNP were between 80 – 100 nm in size, with > 90% encapsulation on mRNA and < 10 EU/mL endotoxin.

### MERS-CoV and SARS-CoV Protein Expression and Purification

Vectors encoding MERS-CoV S-2P^11^ and SARS-CoV S-2P^22^ were generated as previously described with the following small amendments. Proteins were expressed by transfection of plasmids into Expi293 cells using Expifectamine transfection reagent (ThermoFisher) in suspension at 37°C for 4-5 days. Transfected cell culture supernatants were collected, buffer exchanged into 1X PBS, and protein was purified using Strep-Tactin resin (IBA). For proteins used for mouse inoculations, tags were cleaved with addition of HRV3C protease (ThermoFisher) (1% wt/wt) overnight at 4 °C. Size exclusion chromatography using Superose 6 Increase column (GE Healthcare) yielded final purified protein.

### Design and Production of Recombinant Minifibritin Foldon Protein

A mammalian codon-optimized plasmid encoding foldon inserted minifibritin (ADIVLNDLPFVDGPPAEGQSRISWIKNGEEILGADTQYGSEGSMNRPTVSVLRNVEVLDKNIGI LKTSLETANSDIKTIQEAGYIPEAPRDGQAYVRKDGEWVLLSTFLSPALVPRGSHHHHHHSAWS HPQFEK) with a C-terminal thrombin cleavage site, 6x His-tag, and Strep-TagII was synthesized and subcloned into a mammalian expression vector derived from pLEXm. The construct was expressed by transient transfection of Expi293 (ThermoFisher) cells in suspension at 37°C for 5 days. The protein was first purified with a Ni^2+^-nitrilotriacetic acid (NTA) resin (GE Healthcare,) using an elution buffer consisting of 50 mM Tris-HCl, pH 7.5, 400 mM NaCl, and 300 mM imidazole, pH 8.0, followed by purification with StrepTactin resin (IBA) according to the manufacturer’s instructions.

### Cell Lines

HEK293T/17 (ATCC #CRL-11268), Vero E6 (ATCC), Huh7.5 cells (provided by Deborah R. Taylor, US Food and Drug Administration), and ACE-2-expressing 293T cells (provided by Michael Farzan, Scripps Research Institute) were cultured in Dulbecco’s modified Eagle’s medium (DMEM) supplemented with 10% FBS, 2 mM glutamine, and 1% penicillin/streptomycin at 37°C and 5% CO_2_. Vero E6 cells used in plaque assays to determine lung and nasal turbinate viral titers were cultured in DMEM supplemented with 10% Fetal Clone II and 1% anti/anti at 37C and 5% CO2. Vero E6 cells used in PRNT assays were cultured in DMEM supplemented with 10% Fetal Clone II and amphotericin B [0.25 μg/ml] at 37C and 5% CO2. Expi293 cells were maintained in manufacturer’s suggested media.

### *In vitro* mRNA Expression

HEK293T cells were transiently transfected with mRNA encoding SARS-CoV-2 WT S or S-2P protein using a TranIT mRNA transfection kit (Mirus). After 24 hr, the cells were harvested and resuspended in FACS buffer (1X PBS, 3% FBS, 0.05% sodium azide). To detect surface protein expression, the cells were stained with 10 μg/mL ACE2-FLAG (Sigma) or CR3022^30^ in FACS buffer for 30 min on ice. Thereafter, cells were washed twice in FACS buffer and incubated with FITC anti-FLAG (Sigma) or Alexafluor 647 goat anti-human IgG (Southern Biotech) in FACS buffer for 30 min on ice. Live/Dead aqua fixable stain (Invitrogen) were utilized to assess viability. Data acquisition was performed on a BD LSRII Fortessa instrument (BD Biosciences) and analyzed by FlowJo software v10 (Tree Star, Inc.)

### Mouse Models

Animal experiments were carried out in compliance with all pertinent US National Institutes of Health regulations and approval from the Animal Care and Use Committee of the Vaccine Research Center, Moderna Inc., or University of North Carolina at Chapel Hill. For immunogenicity studies, 6-8-week-old female BALB/c (Charles River), BALB/cJ, C57BL/6J, or B6C3F1/J mice (Jackson Laboratory) were used. mRNA formulations were diluted in 50 μL of 1X PBS, and mice were inoculated IM into the same hind leg for both prime and boost. For all SARS-CoV-2 S-P protein vaccinations, mice were inoculated IM, with SAS, as previously detailed^11^. For S + alum immunizations, SARS-CoV-2 S protein (Sino Biological) + 250 μg alum hydrogel was delivered IM. For challenge studies to evaluate MERS-CoV-2 vaccines, 16-20-week-old 288/330^+/+^mice^21^ were immunized. Four weeks post-boost, pre-challenge sera were collected from a subset of mice, and remaining mice were challenged with 5×10^5^ PFU of a mouse-adapted MERS-CoV EMC derivative, m35c4^31^. On day 3 post-challenge, lungs were harvested, and hemorrhage and viral titer were assessed, per previously published methods^32^. For challenge studies to evaluate SARS-CoV-2 vaccines, BALB/cJ mice were challenged with 10^5^ PFU of mouse-adapted SARS-CoV-2 (SARS-CoV-2 MA). On day 2 post-challenge, lungs and nasal turbinates were harvested for viral titer assessment, per previously published methods^28^.

### Histology

Lungs from mice were collected at the indicated study endpoints and placed in 10% neutral buffered formalin (NBF) until adequately fixed. Thereafter, tissues were trimmed to a thickness of 3-5 mm, processed and paraffin embedded. The respective paraffin tissue blocks were sectioned at 5 μm and stained with hematoxylin and eosin (H&E). All sections were examined by a board-certified veterinary pathologist using an Olympus BX51 light microscope and photomicrographs were taken using an Olympus DP73 camera.

### Enzyme-linked Immunosorbent Assay (ELISA)

Nunc Maxisorp ELISA plates (ThermoFisher) were coated with 100 ng/well of protein in 1X PBS at 4°C for 16 hr. Where applicable, to eliminate fold-on-specific binding from MERS S-2P- or SARS-CoV-2 S-2P protein-immune mouse serum, 50 μg/mL of fold-on protein was added for 1 hr at room temperature (RT). After standard washes and blocks, plates were incubated with serial dilutions of heat-inactivated (HI) sera for 1 hr at RT. Following washes, anti-mouse IgG, IgG1, or IgG2a or IgG2c–horseradish peroxidase conjugates (ThermoFisher) were used as secondary Abs, and 3,5,3′5′-tetramethylbenzidine (TMB) (KPL) was used as the substrate to detect Ab responses. Endpoint titers were calculated as the dilution that emitted an optical density exceeding 4X background (secondary Ab alone).

### Pseudovirus Neutralization Assay

We introduced divergent amino acids, as predicted from translated sequences, into the CMV/R-MERS-CoV EMC S (GenBank#: AFS88936) gene^33^ to generate a MERS-CoV m35c4 S gene^31^. To produce SARS-CoV-2 pseudoviruses, a codon-optimized CMV/R-SARS-CoV-2 S (Wuhan-1, Genbank #: MN908947.3) plasmid was constructed. Pseudoviruses were produced by co-transfection of plasmids encoding a luciferase reporter, lentivirus backbone, and S genes into HEK293T/17 cells (ATCC #CRL-11268), as previously described^33^. For SARS-CoV-2 pseudovirus, human transmembrane protease serine 2 (TMPRSS2) plasmid was also co-transfected^34^. Pseudoneutralization assay methods have been previously described^11^. Briefly, HI serum was mixed with pseudoviruses, incubated, and then added to Huh7.5 cells or ACE-2-expressing 293T cells, for MERS-CoV and SARS-CoV-2 respectively. Seventy-two hr later, cells were lysed, and luciferase activity (relative light units, RLU) was measured. Percent neutralization was normalized considering uninfected cells as 100% neutralization and cells infected with only pseudovirus as 0% neutralization. IC_50_ titers were determined using a log (agonist) vs. normalized response (variable slope) nonlinear function in Prism v8 (GraphPad).

### Plaque Reduction Neutralization Test (PRNT)

HI sera were diluted in gelatin saline (0.3% [wt/vol] gelatin in phosphate-buffered saline supplemented with CaCl_2_ and MgCl_2_) to generate a 1:5 dilution of the original specimen, which served as a starting concentration for further serial log_4_ dilutions terminating in 1:81,920. Sera were combined with an equal volume of SARS-CoV-2 clinical isolate 2019-nCoV/USA-WA1-F6/2020 in gelatin saline, resulting in an average concentration of 730 plaque-forming units per mL (determined from plaque counts of 24 individual wells of untreated virus) in each serum dilution. Thus, final serum concentrations ranged from 1:10 to 1:163,840 of the original. Virus/serum mixtures were incubated for 20 min at 37 °C, followed by adsorption of 0.1 mL to each of two confluent Vero E6 cell monolayers (in 10-cm^2^ wells) for 30 min at 37°C. Cell monolayers were overlaid with Dulbecco’s modified Eagle’s medium (DMEM) containing 1% agar and incubated for 3 d at 37°C in humidified 5% CO_2_. Plaques were enumerated by direct visualization. The average number of plaques in virus/serum (duplicate) and virus-only (24) wells was used to generate percent neutralization curves according the following formula: 1 – (ratio of mean number of plaques in the presence and absence of serum). The PRNT IC_50_ titer was defined as the reciprocal serum dilution at which the neutralization curve crossed the 50% threshold.

### Intracellular Cytokine Staining

Mononuclear single cell suspensions from whole mouse spleens were generated using a gentleMACS tissue dissociator (Miltenyi Biotec) followed by 70 μm filtration and density gradient centrifugation using Fico/Lite-LM medium (Atlanta Biologicals). Cells from each mouse were resuspended in R10 media (RPMI 1640 supplemented with Pen-Strep antibiotic, 10% HI-FBS, Glutamax, and HEPES) and incubated for 6 hr at 37°C with protein transport inhibitor cocktail (eBioscience) under three conditions: no peptide stimulation, and stimulation with two spike peptide pools (JPT product PM-WCPV-S-1). Peptide pools were used at a final concentration of 2 μg/mL each peptide. Cells from each group were pooled for stimulation with cell stimulation cocktail (eBioscience) as a positive control. Following stimulation, cells were washed with PBS prior to staining with LIVE/DEAD Fixable Blue Dead Cell Stain (Invitrogen) for 20 min at RT. Cells were then washed in FC buffer (PBS supplemented with 2% HI-FBS and 0.05% NaN_3_) and resuspended in BD Fc Block (clone 2.4G2) for 5 min at RT prior to staining with a surface stain cocktail containing the following antibodies purchased from BD and Biolegend: I-A/I-E (M5/114.15.2) PE, CD8a (53-6.7) BUV805, CD44 (IM7) BUV395, CD62L (MEL-14) BV605, and CD4 (RM4-5) BV480 in brilliant stain buffer (BD). After 15 min, cells were washed with FC buffer then fixed and permeabilized using the BD Cytofix/Cytoperm fixation/permeabilization solution kit according to manufacturer instructions. Cells were washed in perm/wash solution and stained with Fc Block (5 min at RT), followed by intracellular staining (30 min at 4°C) using a cocktail of the following antibodies purchased from BD, Biolegend, or eBioscience: CD3e (17A2) BUV737, IFN-γ (XMG1.2) BV650, TNF-α (MP6-XT22) BV711, IL-2 (JES6-5H4) BV421, IL-4 (11B11) Alexa Fluor 488, and IL-5 (TRFK5) APC in 1x perm/wash diluted with brilliant stain buffer. Finally, cells were washed in perm/wash solution and resuspended in 0.5% PFA-FC stain buffer prior to running on a Symphony A5 flow cytometer (BD). Analysis was performed using FlowJo software, version 10.6.2 according to the gating strategy outlined in **Extended Data Figure 9**. Background cytokine expression in the no peptide condition was subtracted from that measured in the S1 and S2 peptide pools for each individual mouse.

### T Cell Stimulation and Cytokine Analysis

Spleens from immunized mice were collected 2 weeks post-boost. 2 × 10^6^ splenocytes/well (96-well plate) were stimulated *in vitro* with two peptide libraries, JPT1 and JPT2, (15mers with 11 aa overlap) covering the entire SARS-CoV-2 spike protein (JPT product PM-WCPV-S-1). Both peptide libraries were used at a final concentration of 1 μg/mL. After 24 hr of culture at 37°C, the plates were centrifuged and supernatant was collected and frozen at −80°C for cytokine detection. Measurements and analyses of secreted cytokines from a murine 35-plex kit were performed using a multiplex bead-based technology (Luminex) assay with a Bio-Plex 200 instrument (Bio-Rad) after 2-fold dilution of supernatants.

### Statistical Analysis

Geometric means or means are represented by the heights of bars, or symbols, and error bars represent the corresponding SD. Dotted lines indicate assay limits of detection. Mann-Whitney tests were used to compare 2 experimental groups and Wilcoxon signed rank tests to compare the same animals at different time points. To compare >2 experimental groups, Kruskal-Wallis ANOVA with Dunn’s multiple comparisons tests were applied. For antibody responses in **Extended Data Fig. 4c**, a Spearman correlation test was used to correlate binding antibody titers to neutralizing antibody titers. * = p-value < 0.05, ** = p-value < 0.01, *** = p-value < 0.001, **** = p-value < 0.0001.

---

**Extended Data Figure 1.**
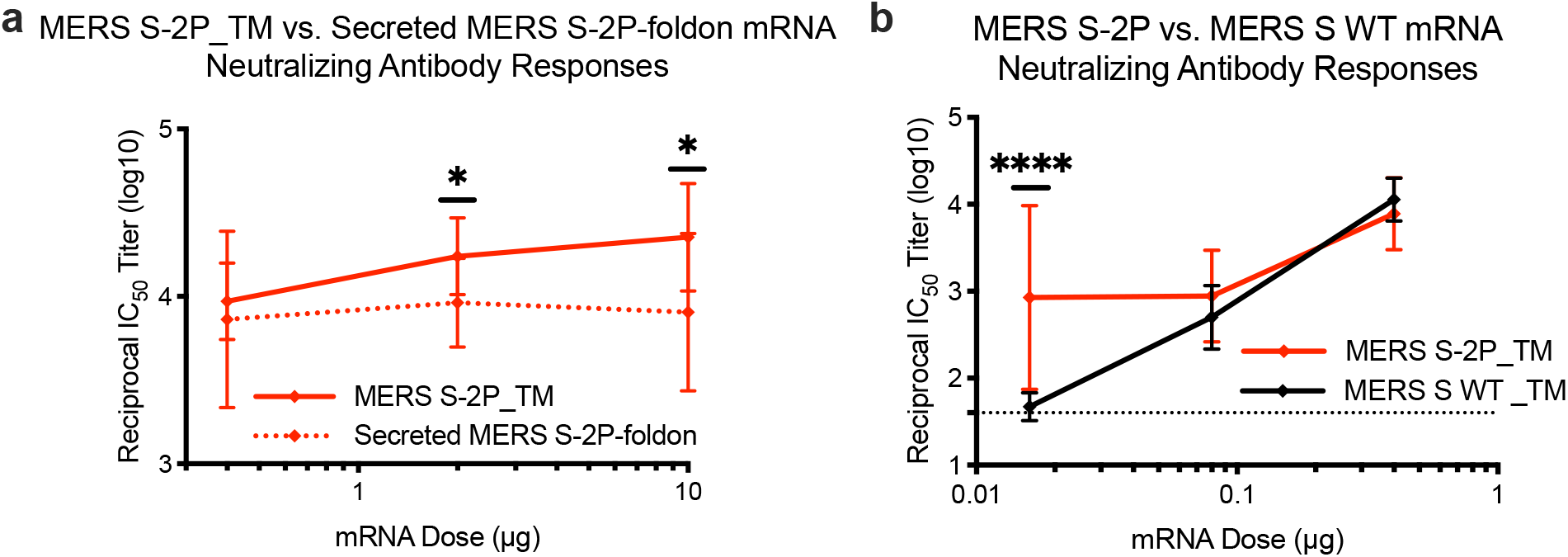
Transmembrane-anchored MERS-CoV S-2P (S-2P_TM) mRNA elicits more potent neutralizing antibody responses than secreted MERS-CoV S-2P and S WT mRNA. C57BL/6J mice were immunized at weeks 0 and 4 with (a) 0.4, 2, or 10 μg of MERS-CoV S-2P_TM (red) or MERS S-2P_secreted (red hashed) or (b) 0.016 μg, 0.08 μg, or 0.4 μg of MERS-CoV S-2P or MERS-CoV S WT_TM (black) mRNA. Sera were collected 4 weeks post-boost and assessed for neutralizing antibodies against MERS-CoV m35c4 pseudovirus. Dotted line = assay limit of detection. Immunogens were compared at each dose level

**Extended Data Figure 2.**
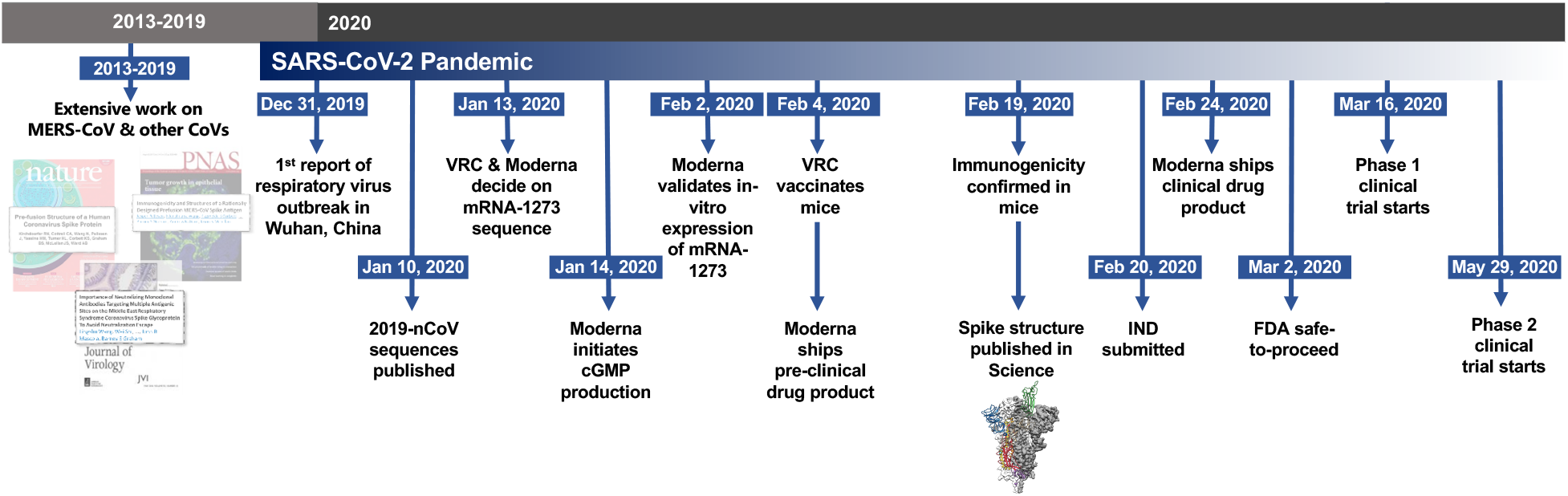
Timeline for mRNA-1273’s progression to clinical trial. The morning after novel coronavirus (nCoV) sequences were released, spike sequences were modified to include prefusion stabilizing mutations and synthesized for protein production, assay development, and vaccine development. Twenty-five days after viral sequences were released, clinically-relevant mRNA-1273 was received to initiate animal experiments. Immunogenicity in mice was confirmed 15 days later. Moderna shipped clinical drug product 41 days after GMP production began, leading to the Phase 1 clinical trial starting 66 days following the release of nCoV sequences.

**Extended Data Figure 3.**
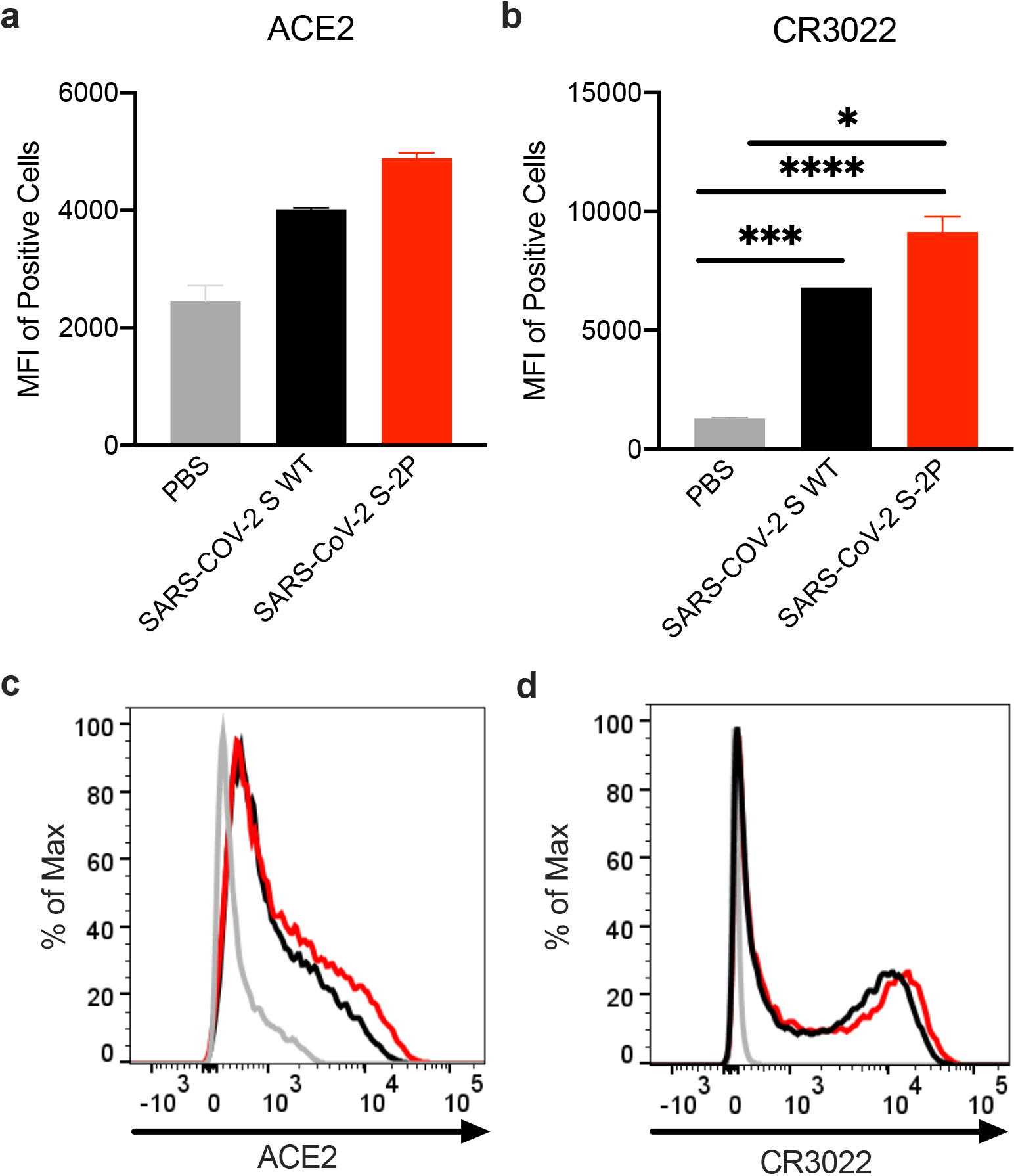
*In vitro* expression of SARS-CoV-2 spike mRNA on cell surface. 293T cells were transfected with mRNA expressing SARS-CoV-2 wild-type spike (black) or S-2P (red), stained with ACE2 (a,c) or CR3022 (b,d), and evaluated by flow cytometry 24 post-transfection. Mock-transfected (PBS) cells served as a control.

**Extended Data Figure 4.**
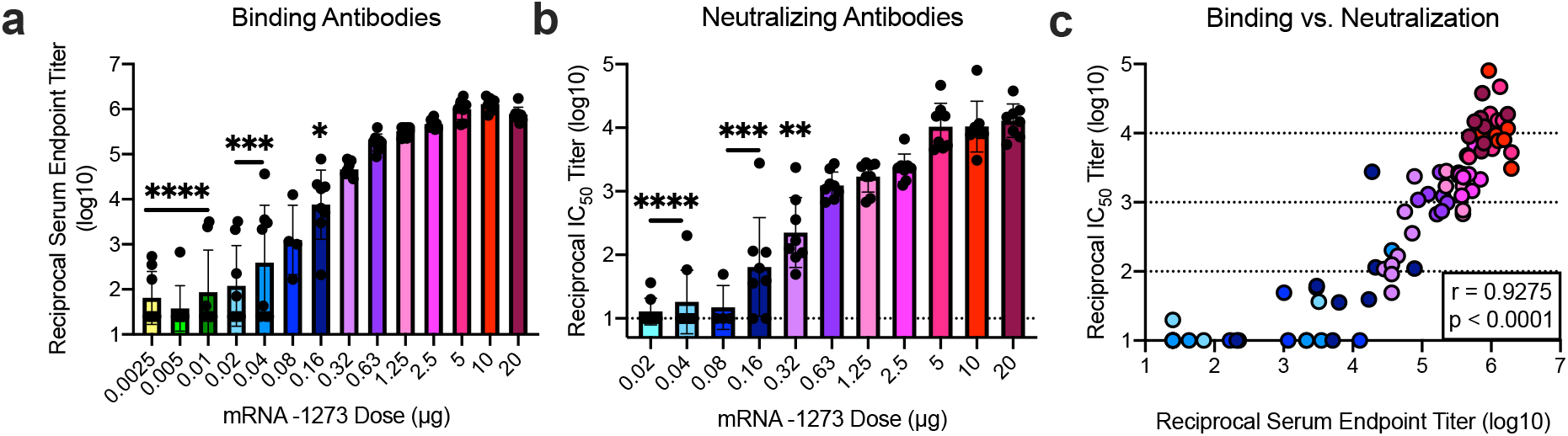
Dose-dependent mRNA-1273-elicited antibody responses reveal strong positive correlation between binding and neutralization titers. BALB/cJ mice were immunized at weeks 0 and 3 weeks with various doses (0.0025 − 20 μg) of mRNA-1273. (a-b) Sera were collected 2 weeks post-boost and assessed for SARS-CoV-2 S-specific IgG by ELISA (a) and neutralizing antibodies against homotypic SARS-CoV-2 pseudovirus (b). (a-b) All doses were compared to 20 μg dose.

**Extended Data Figure 5.**
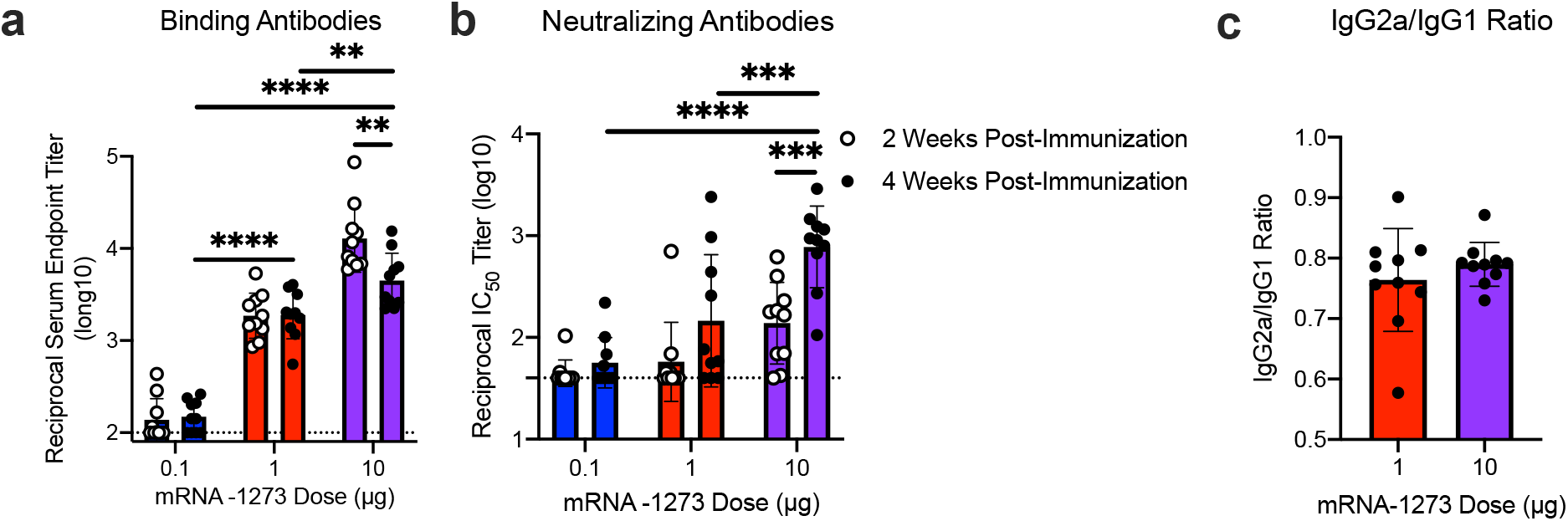
A single dose of mRNA-1273 elicits robust antibody responses. BALB/cJ mice were immunized with 0.1 (blue), 1 μg (red), or 10 μg (purple) of mRNA-1273. Sera were collected 2 (open circles) and 4 (closed circles) weeks post-immunization and assessed for SARS-CoV-2 S-specific total IgG by ELISA (a) and neutralizing antibodies against homotypic SARS-CoV-2 pseudovirus (b). (c) S-specific IgG2a and IgG1 were also measured by ELISA, and IgG2a to IgG1 subclass ratios were calculated. Dotted line = assay limit of detection. (a-b) Doses were compared 4 weeks post-boost, and timepoints were compared within each dose level.

**Extended Data Figure 6.**
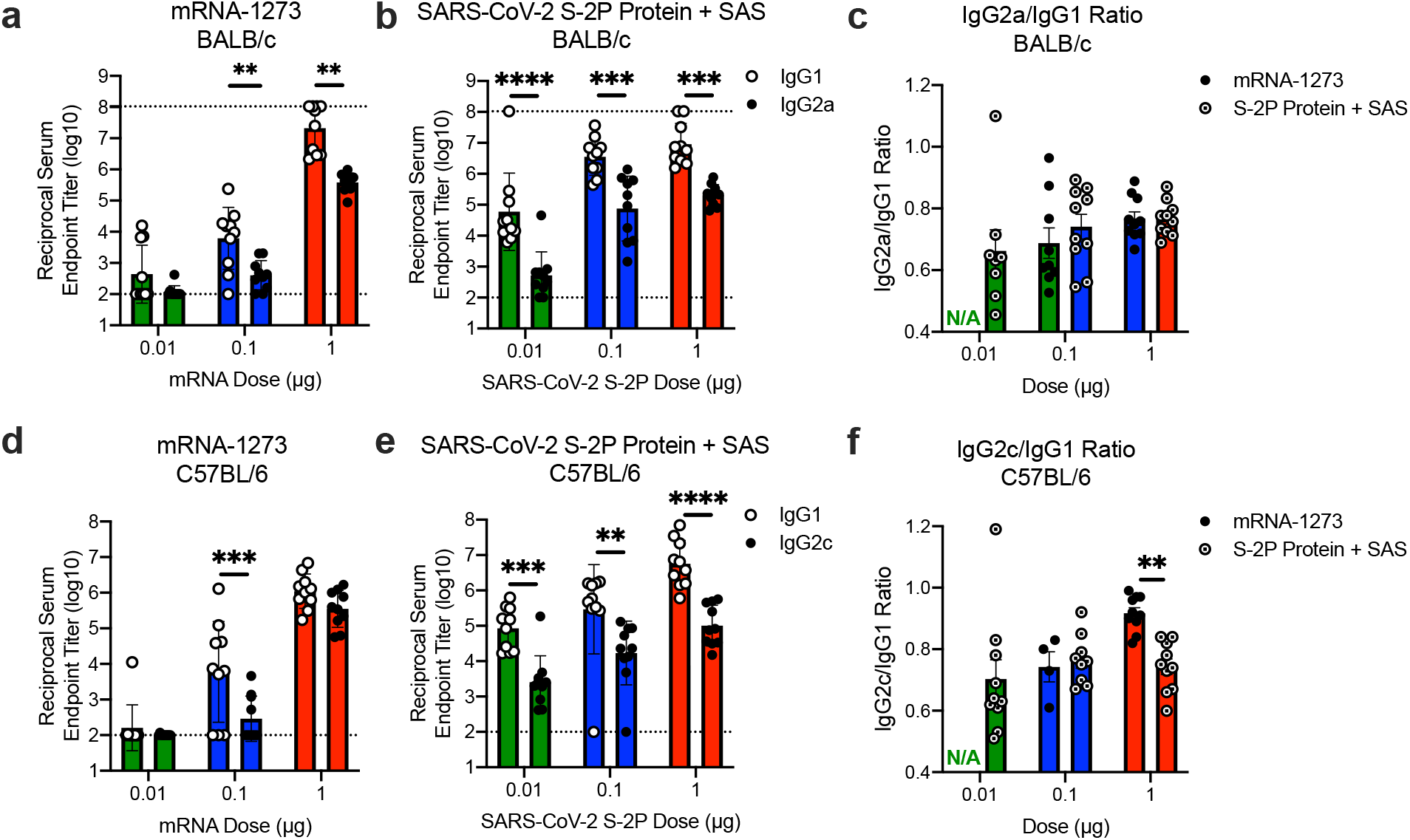
mRNA-1273 and SAS-adjuvanted S-2P protein elicit both IgG2a and IgG1 subclass S-binding antibodies. BALB/cJ (a-c) or C57BL/6J (d-f) mice were immunized at weeks 0 and 3 with 0.01 (green), 0.1 (blue), or 1 μg (red) of mRNA-1273 SARS-CoV-2 S-2P protein adjuvanted with SAS. Sera were collected 2 weeks post-boost and assessed by ELISA for SARS-CoV-2 S-specific IgG1 and IgG2a or IgG2c for BALB/cJ and C57BL/6J mice, respectively. Endpoint titers (a-b, d-e) and endpoint titer ratios of IgG2a to IgG1 (c) and IgG2c to IgG1 (f) were calculated. For mice for which endpoint titers did not reach the lower limit of detection (dotted line), ratios were not calculated (N/A). IgG1 and IgG2a/c (a-b, d-e) and immunogens (c, f) were compared at each dose level.

**Extended Data Figure 7.**
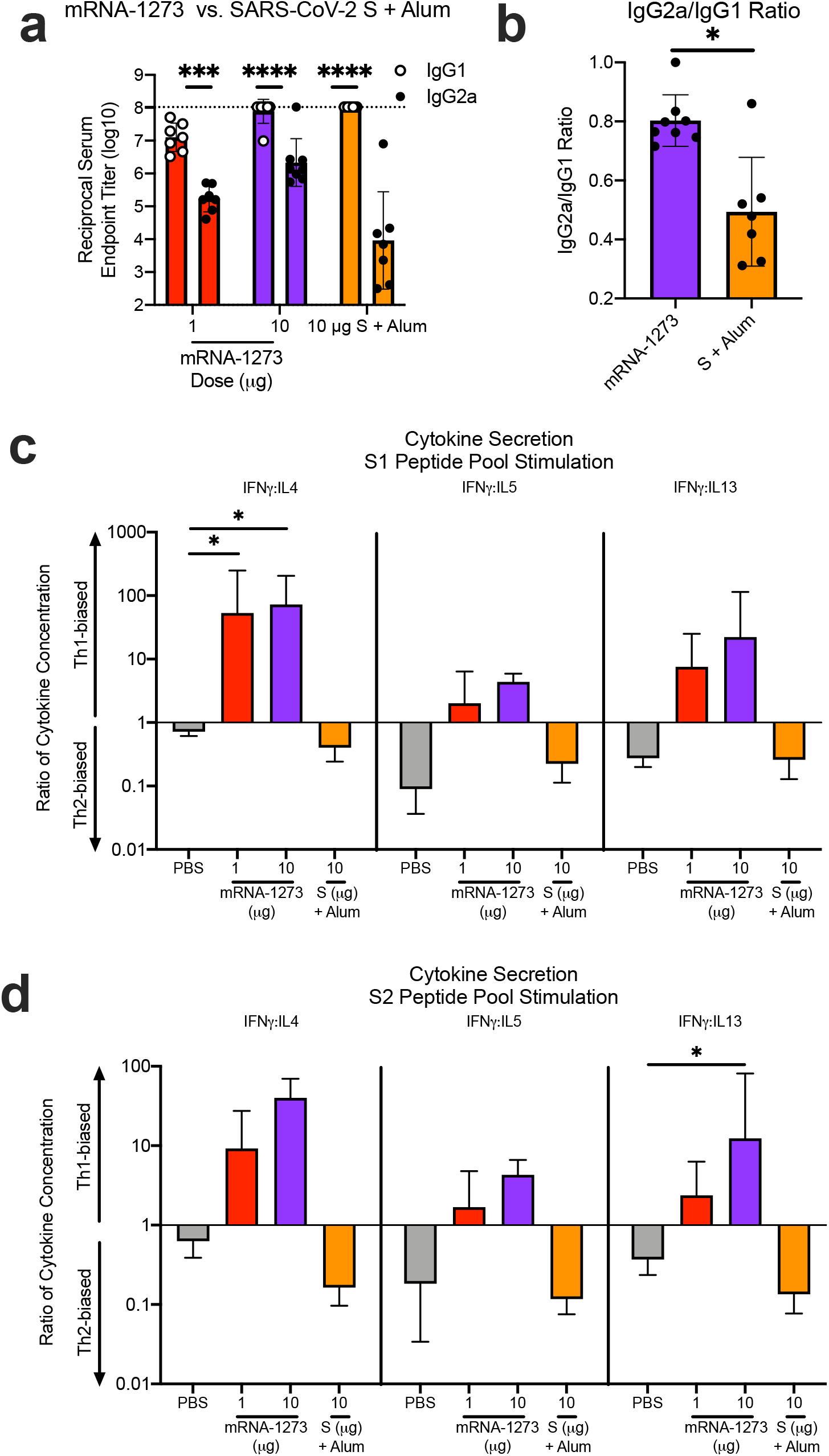
mRNA-1273 elicits Th1-skewed responses compared to S protein adjuvanted with alum. BALB/c mice were immunized at weeks 0 and 2 weeks with 1 (red) or 10 μg (purple) of mRNA-1273 or 10 μg of SARS-CoV-2 S protein adjuvanted with alum hydrogel (orange). (a-b) Sera were collected 2 weeks post-boost and assessed by ELISA for SARS-CoV-2 S-specific IgG1 and IgG2a. Endpoint titers (a) and endpoint titer ratios of IgG2a to IgG1 (b) were calculated. (c-d) Splenocytes were also collected 4 weeks post-boost to evaluate IFN-γ IL-4, IL-5, and IL-13 cytokine levels secreted by T cells re-stimulated with S1 (c) and S2 (d) peptide pools, measured by Luminex. Dotted line = assay limit of detection. IgG1 and IgG2a/c (a) were compared at each dose level. (c-d) For cytokines, all comparisons were compared to PBS-immunized mice.

**Extended Data Figure 8.**
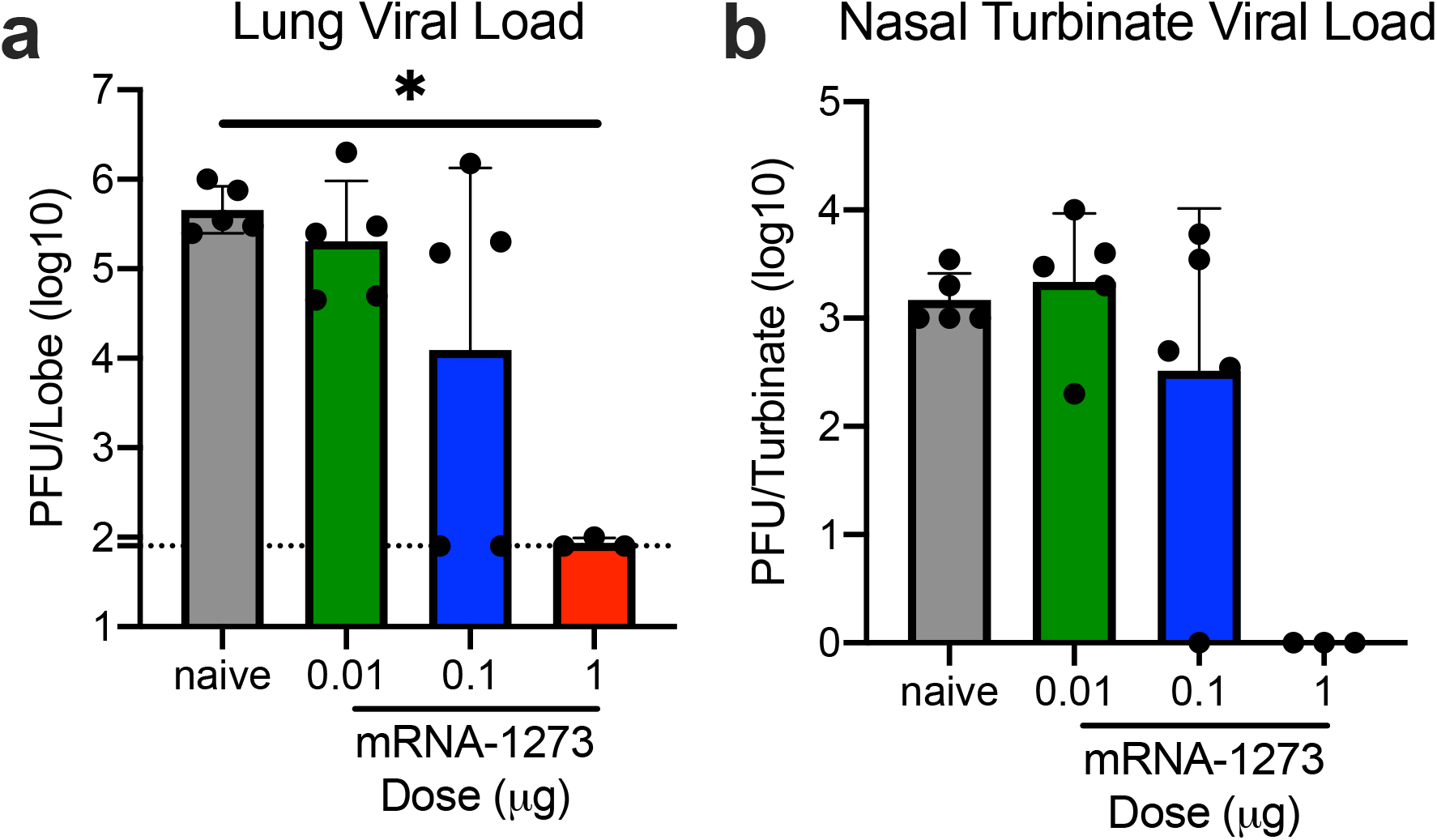
mRNA-1273 protects mice from upper and lower airway SARS-CoV-2 infection, 13 weeks post-boost. BALB/cJ mice were immunized at weeks 0 and 3 with 0.01 (green), 0.1 (blue), or 1 μg (red) of mRNA-1273. Age-matched naive mice (gray) served as controls. Thirteen weeks post-boost, mice were challenged with mouse-adapted SARS-CoV-2. Two days post-challenge, at peak viral load, mouse lungs (a) and nasal turbinates (b) were harvested from 5 mice per group for analysis of viral titers. Dotted line = assay limit of detection. All dose levels were compared.

**Extended Data Figure 9.**
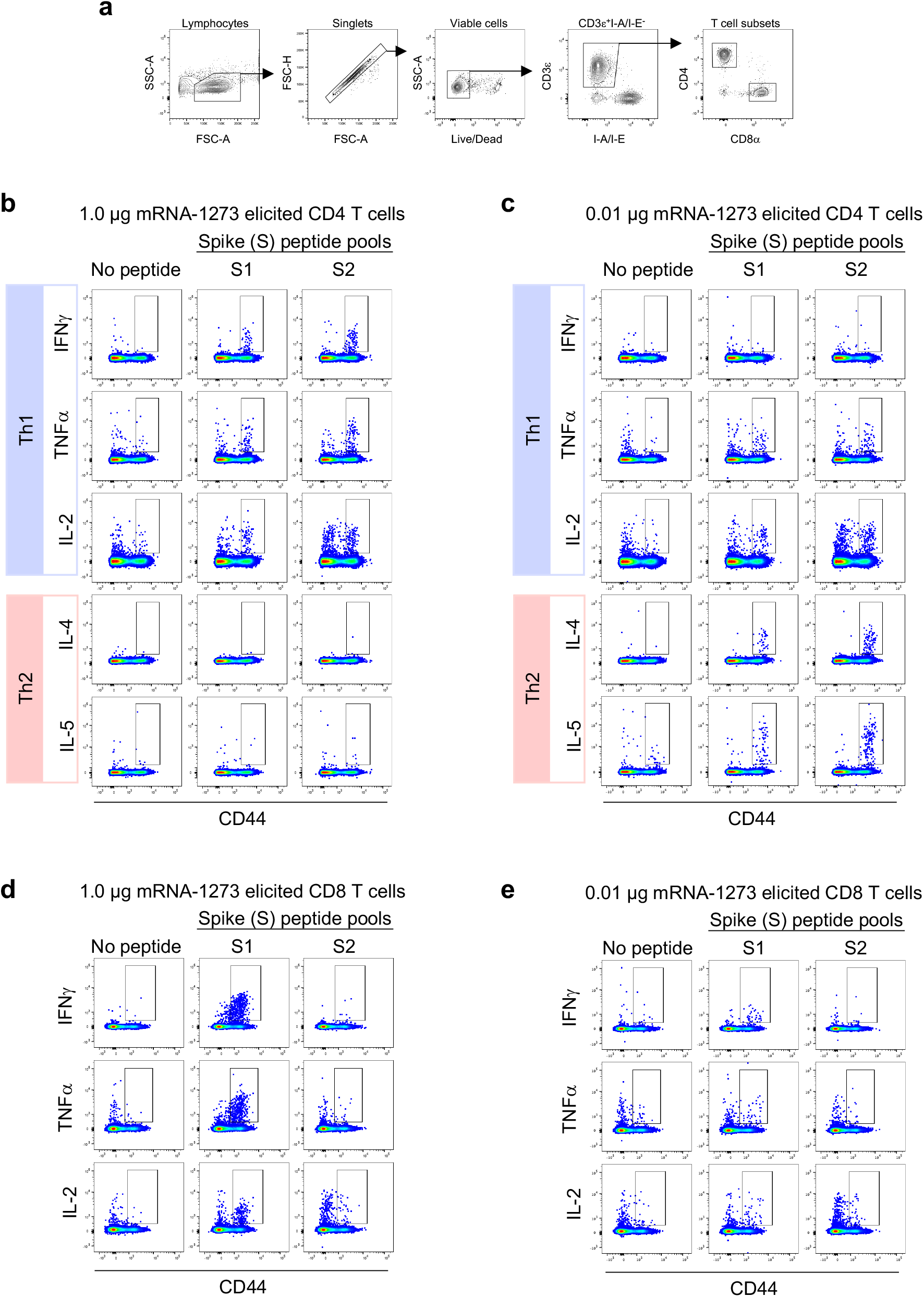
Flow cytometry panel to quantify SARS-CoV-2 S-specific T cells in mice. (a) A hierarchical gating strategy was used to unambiguously identify single, viable CD4+ and CD8+ T cells. Gating summary of SARS-CoV-2 S-specific (b-c) CD4 (b-c) and (d-e) CD8 (d-e) T cells elicited by 1.0 and 0.01 μg mRNA-1273 immunization. Antigen-specific T cell responses following peptide pool re-stimulation were defined as CD44^hi^/cytokine^+^. Concatenated files shown were generated using the same number of randomly selected events from each animal across the different stimulation conditions using FlowJo software, v1

**Extended Data Table 1.** Concordance of Pseudovirus Neutralization Assay and PRNT.

| Mouse Serum<br>Pool #<sup>1</sup> | Reciprocal IC<sub>50</sub> Titer |  | *Fold*<br>*Difference*<sup>4</sup> |
| --- | --- | --- | --- |
|  | Pseudovirus Neutralization<sup>2</sup> | PRNT<sup>3</sup> |  |
| 1 | 893.5 +/- 1.4 | 933.5 | 1.0 |
| 2 | 211.6 +/- 1.5 | 314.5 | 0.7 |
| 3 | 159.8 +/- 1.3 | 397.1 | 0.5 |

<sup>1</sup>BALB/cJ mice were immunized at weeks 0 and 3 with 1 µg SARS-CoV-2 S-2P protein, adjuvanted with SAS. Sera were collected 2 weeks post-boost and pooled (N = 3 mice/pool).
<sup>2</sup>IC<sub>50</sub> titers were averaged from pseudovirus neutralization assays completed in 5 experimental replicates. (GMT +/- geometric SD)
<sup>3</sup>IC<sub>50</sub> titer from PRNT assay completed once.
<sup>4</sup>Fold difference calculated as average pseudovirus neutralization IC<sub>50</sub> titer relative to PRNT IC<sub>50</sub> titer.

